# Methionine and Iron Supplementation Enhances Kanamycin Activity Against *Mycobacterium tuberculosis*

**DOI:** 10.1101/2023.02.28.530446

**Authors:** Hanumantharao Paritala, Kate S. Carroll

## Abstract

Despite understanding the functional roles of important proteins in sulfate assimilation and trans-sulfuration pathways, no study has been conducted to estimate the changes in the intracellular thiol levels when mycobacterium is treated with methionine. Methionine was known to be sulfur source for mycobacteria when it resides in the hostile granuloma environment and through transsufuration pathway methionine was shown to stimulate the sulfate assimilation. To study the dynamics of sulfate assimilation pathway we created gene knockout strains of important enzymes in the sulfate assimilation pathway of M. smegmatis. We stimulated these mutants with methionine, pro-filed the changes in intracellular thiols, applied external oxidative stress, and monitored the associated intracellular redox potentials (EMSH). These experiments revealed that when M. smegmatis cultures are supplemented with methionine intracellular levels surged. When those cells were challenged with hydrogen peroxide the M. smegmatis cells showed increased intracellular oxidative potentials and took prolonged time to revert back normal redox balance state when compared to non-methionine containing controls. Here in this report, we are describe how the dynamic changes of intracellular thiols are linked to augment the activity of kanamycin.

## Introduction

Tuberculosis (TB), is an infectious bacterial disease caused by Mycobacterium tuberculosis. It is transmitted from person to person via inhalation of aerosol droplets from the throat and lungs of people with the active respiratory disease. TB can be treated by taking several first line drugs for 6 to 9 months. Improper administration of treatment regimens and failure to ensure that drug-susceptible TB patients complete the whole course of treatment in chemotherapy results in drug resistance. TB which is resistant to at least two frontline drugs, rifampin and isoniazid, is considered as multidrug resistant tuberculosis (MDR-TB). However recent studies consider that TB resistant to the fluoroquinolones and streptomycin but susceptible to second-line injectables, kanamycin, amikacin and capreomycin, is also treated as MDR-TB (1). Treatment of MDR-TB is challenging because of the high toxicity of second-line drugs, a smaller number of treatment options and the longer treatment duration required compared with drug-susceptible TB. The selection of drugs in MDR-TB is based on previous treatment history, drug susceptibility results, and TB drug resistance patterns in the each region (2). The kanamycin is the initial choice of injectable durgs and followed by amikacin, capreomycin and newer fluoroquinolones include levofloxacin and moxifloxacin as recommended by World Health Organization (3). Mismanagement of MDR-TB allows the bacilli to advance to extensively drug-resistant (XDR) variety which is resistant to treatment with any fluoroquinolone and any of the injectable drugs used in treatment (kanamycin, amikacin and capreomycin). MDR tuberculosis and XDR tuberculosis are serious threats to the progress that has been made in the control of tuberculosis worldwide over the past decade (4,5). The second line drugs kanamycin and amikacin are aminoglycoside analogs and share hundred percent of cross resistance to each other and do not share with streptomycin. Hence containing and management of MDR-TB is critical and important. Any improvement or potentiation in the activity of the second line antibiotics, kanamycin/amikacyn is greatly beneficial for controlling the worldwide tuberculosis spread of resistant tuberculosis and decreases the cost and duration of the treatment.

Previous researchers have attempted to potentiate antibiotics by several strategies which are specific to given condition and antibiotic in use. Lebeaux *et al* demonstrated that a combination of gentamicin and the clinically compatible basic amino acid L-arginine increased the in vitro planktonic and biofilm susceptibility to gentamicin, with 99% mortality amongst clinically relevant pathogens, i.e. *S. aureus, E. coli* and *P. aeruginosa* persistent bacteria. (6), beta-lactam antibitics get potentiated by co administering with beta-lactamase inhibitor calvulanate (7,8) and Trimethoprim is commonly co-administered with sulfonamides, for example sulfamethoxazole in the co-trimoxazole combination, to achieve synergy (9,10) in preventing bacterial infections. As the rate of drug resistance expansion appears far beyond that of the current drug developmental process, it is anticipated that such augmentation approaches will work to improve the utility and protection of those effective agents. In this report we are describing one such approach which utilizes the dynamics of sulfate assimilation pathway in mycobacteria to improve the activity of kanamycin.

Despite understanding the functional roles of important proteins in sulfate assimilation and transsulfuration pathways, no study has been conducted to estimate the changes in the intracellular thiol levels when mycobacterium is treated with methionine. Methionine was known to be sulfur source for mycobacteria when it resides in the hostile granuloma environment and through transsufuration pathway methionine was shown to stimulate the sulfate assimilation (11). To study the dynamics of sulfate assimilation pathway we created gene knockout strains of important enzymes in the sulfate assimilation pathway of *M. smegmatis*. We stimulated these mutants with methionine, profiled the changes in intracellular thiols, applied external oxidative stress, and monitored the associated intracellular red-ox potentials (*E*_*MSH*_). These experiments revealed that when *M. smegmatis* cultures are supplemented with methionine intracellular levels surged. When those cells were challenged with hydrogen peroxide the *M. smegmatis* cells showed increased intracellular oxidative potentials and took prolonged time to revert back normal red-ox balance state when compared to non-methionine containing controls. Here in this report, we are describing how the dynamic changes of intracellular thiols were linked to augment the activity of kanamycin.

## Results

### E_MSH_ measurement in M. Smegmatis sulfate assimilation pathway knock out strains

*Cys*A mutant exhibited oxidized *E*_*MSH*_ while the *cys*H mutant displayed reduced *E*_*MSH*_ as compared to wt *M. smegmatis*. However cysH mutant was unable to grow in the absence of methionine and hence the E_MSH_. Addition of exogenous methionine restored the intra bacterial *E*_*MSH*_ values comparable to the wt in both the mutants. The *E*_*MSH*_ of the *cys*Q remained unaffected and was identical to the wt *M. smegmatis*. Consistent with role of Mycothionine reductase, the ΔMtr mutant exhibited oxidized *E*_*MSH*_ that showed no significant change in *E*_*MSH*_ upon addition of methionine.

### E_MSH_ measurement in M. Smegmatis sulfate assimilation pathway knock out strains in presence of hydrogenperoxide

The cultures were grown in presence and absence of 0.3 mM methionine and exposed to 500μM H_2_O_2_. We observed that oxidative stress induces similar degree of roGFP2 oxidation in wt, cysA, and cysQ strains except in Δmtr strain as indicated by a comparable increase in the 405/488 ratio Figure 2a. However, a striking difference was evident during the recovery phase. The Δmtr mutant showed extended oxidation (or delayed recovery), followed by cysA, cysQ and wt *M smegmatis*. Although this was expected in the case of cysA mutant because of the oxidized steady-state *E*_*MSH*_ observed (Figure 1), and in Δmtr due to the inability of the strain to reduce mycothione. However delayed recovery was surprising in the case of wt *M. smegmatis* and cysQ mutants. Next, we analyzed the effect of exogenous methionine on the dynamic response. Interestingly, we observed that addition of methionine significantly prolonged the oxidative burst in all the strains. However, in comparison to wt *M.smegmatis*, the effect was moderate in the case of Δmtr and cysH:Q mutants and substantial in the case of the cysA mutant and we were unable to observe any recovery from oxidative stress in the cysA+ methionine mutants during the 30 min time course of this assay. We think that addition of methionine may altered the homeostasis of thiol pool of the cells, possibly leading to increased production of cysteine that may lead to excessive oxidative stress and therefore a prolonged oxidative burst observed with methionine treated cells. Park and Imlay (2003) have similarly reported that addition of cystine followed by H_2_O_2_treatment perturbs the total thiol content in *E. coli*, particularly the cysteine content of the cell, which led to fueling of the Fenton reaction(12).

**Figure 1.**
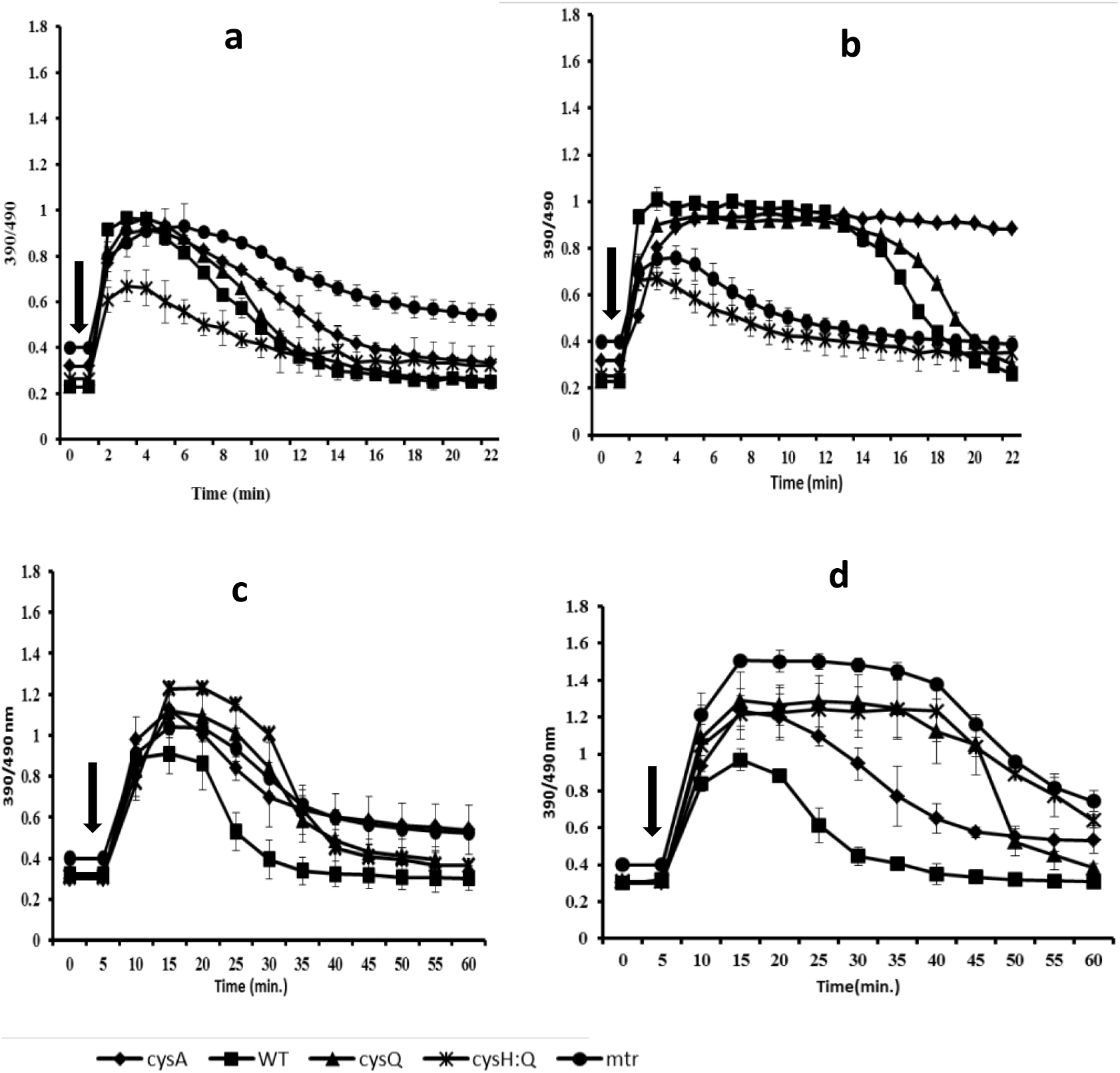
Dynamic response of Mrx1-roGFP2 bearing strains of mycobacteria. *M. smegmatis* and mutant strains exposed to 500μM H_2_O_2_ (a) & (b) or 50 μM CHP (c & d) a and c represent the response in absence of exogenous methionine while (b and d) in presence of 0.3mM methionine.

**Figure 2:**
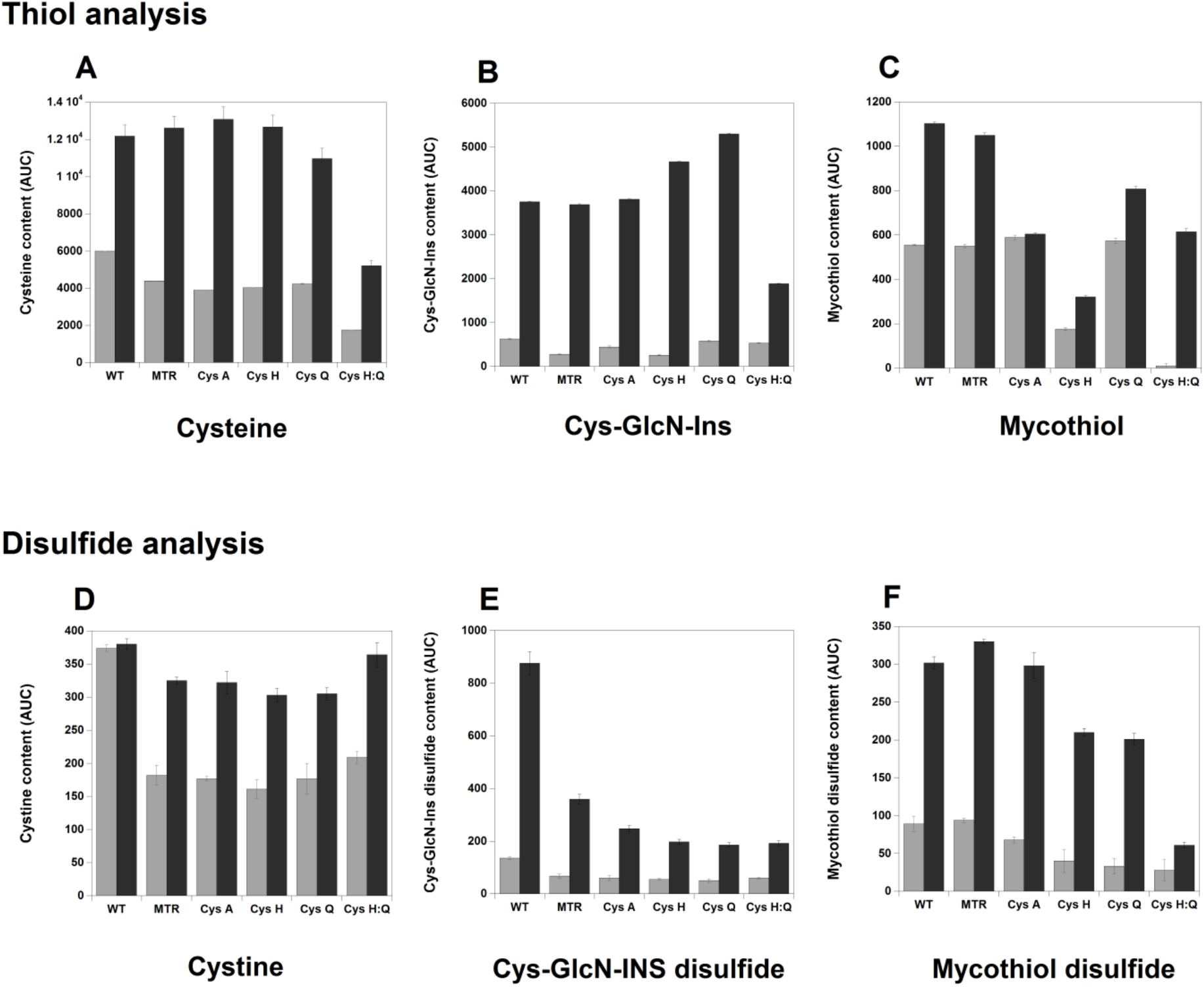
Mycothiol and its components analysis of *M. Smegmatis* knockout strains in presence and in absence of methionine.

### Thiol analysis of M. Smegmatis knockout strains in presence and in absence of Methionine

Mycothiol and its components, cysteine, cys-glcn-ins contents in *M.smegmatis* and gene knockout strains were found to be elevated 2 to 3 fold when the cultures grown in presence of methionine. Analysis of the disulfide content also revealed 1 to 5 fold increase in oxidation of the thiols. The facile autoxidation of cysteine has been attributed to the presence of a free amino group located beta to the thiol group that facilitates binding of heavy metals to catalyze thiol autoxidation. cys-glc-ins also possesses this functionality. In formyl-Cys-GlcN-Ins, succ-Cys-GlcN-Ins, and MSH, the amino group is acylated so that autoxidation is expected to be much slower. Accordingly, we observe relatively low levels of mycothiol disulfide despite being a dominant thiol in *M. smegmatis*. The relatively oxidized state of Cys-GlcN-Ins might result from Cys-GlcN-Ins undergoing autoxidation in the cell at an unusually rapid rate or from lack of an adequate system to reduce its oxidized products. Experiments conducted by Newton et al revealed that the initial rate (0 to 8 min) of oxidation of Cys-GlcN Ins was ∼11-fold faster than that of MSH when tested independently at 200 μM initial thiol concentrations and copper as the catalyst(13). A positively charged ammonium residue, in the case of CysGlcN-Ins, might render the molecule as a poor substrate for Mycothione reductase (MTR) to get converted to the reduced form. Also, the positive charge may facilitate the binding of heavy metals like copper, iron to form in to disulfide(13). Taken together these experiments demonstrate that treatment of *M. smegmatis* cells with methionine resulted in the elevation of intracellular thiol levels, particularly cysteine, cys-glcN-Ins and Mycothiol. Accumulation of these thiols, together with the available iron, could lead to generation of unwanted oxidative stress through Fenton type reaction. If this is the case, coupling of this oxidative stress with the activity of anti-tubercular drugs should result in enhanced activity of the antibiotics.

### *M. smegmatis killing experiments in presence of methionine and iron* potentiated the kanamycin activity

Kanamycin was tested against *M. smegmatis* in vitro and was consistently observed to be potentiated by 10 fold when co administered along with methionine and iron. This indicates that the methionine induced cellular thiol levels might be undergoing Fenton type reaction in presence of iron and are elevating the intracellular oxidative stress. In presence of kanamycin the generated oxidative stress might be synergized with antibiotic activity and might have resulted in the potentiation of antibiotic activity.

### QPCR experiments to quantify the extent of DNA damage occurred for M. smegmatis when treated with antibiotics in presence of methionine and iron

The vulnerability of cells to oxidative DNA damage is strongly affected by their physiological state. The methionine treatment causes disruption in cysteine homeostasis and results in elevation of cysteine and other thiol levels. In vitro analysis demonstrated that cysteine reduces ferric iron with exceptional speed and this action permits free iron to redox cycle rapidly in presence of hydrogen peroxide and increases the rate of hydroxy radicals formation. During normal growth conditions cells maintain small cysteine and other thiol levels and thus thiols are not major contributors for DNA damage (12). In line with the previous observations we found that methionine and iron treatment up-regulated the cysteine and other thiol levels in *M. smegmatis* and when challenged these cells with kanamycin in presence of iron we found 2 fold increase in DNA damage when compared to kanamycin only treated *M. smegmatis*.

### Effect of exogenous methionine and Fe on kanamycin toxicity in *M. tuberculosis* H37Rv

Exogenous methionine and Fe and methionine alone potentiated the effect of kanamycin as can be observed with reduced survival in case of iron + methionine and methionine alone (Figure 3b). Two fold decrease in MIC with respect to control was observed when cells pretreated with methionine and Fe + methionine were plated on different concentration of kanamycin (Fig. 3b). The Mrx1-roGFP sensor response was also checked upon treatment with kanamycin and is consistent with the observation in *M.smegmatis* kanamycin was able to elicit a strong oxidative sensor response in presence of methionine and Fe + methionine. Vaubourgeix et al., (2015) demonstrated the presence irreversibly oxidized proteins, a common marker for oxidative stress, upon kanamycin treatment in *Mtb* and *M.smegmatis* which further supports the involvement of oxidative stress in kanamycin activity (14).

**Figure 3:**
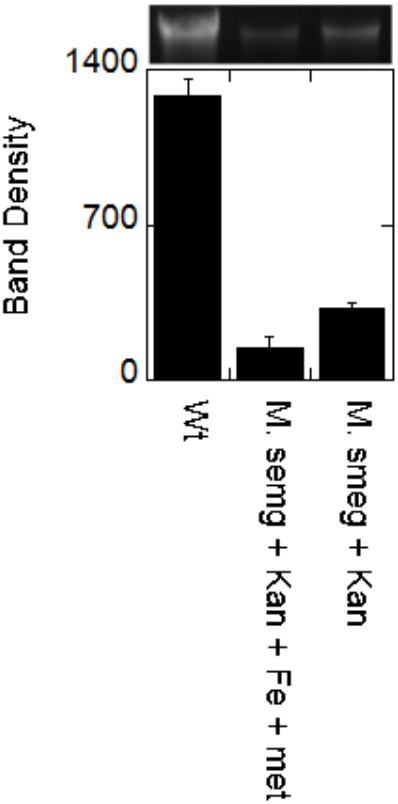
QPCR experiments to quantify the extent of DNA damage occurred for M. smegmatis when treated with Kanamycin in presence and absence of methionine and iron.

**Figure 4.**
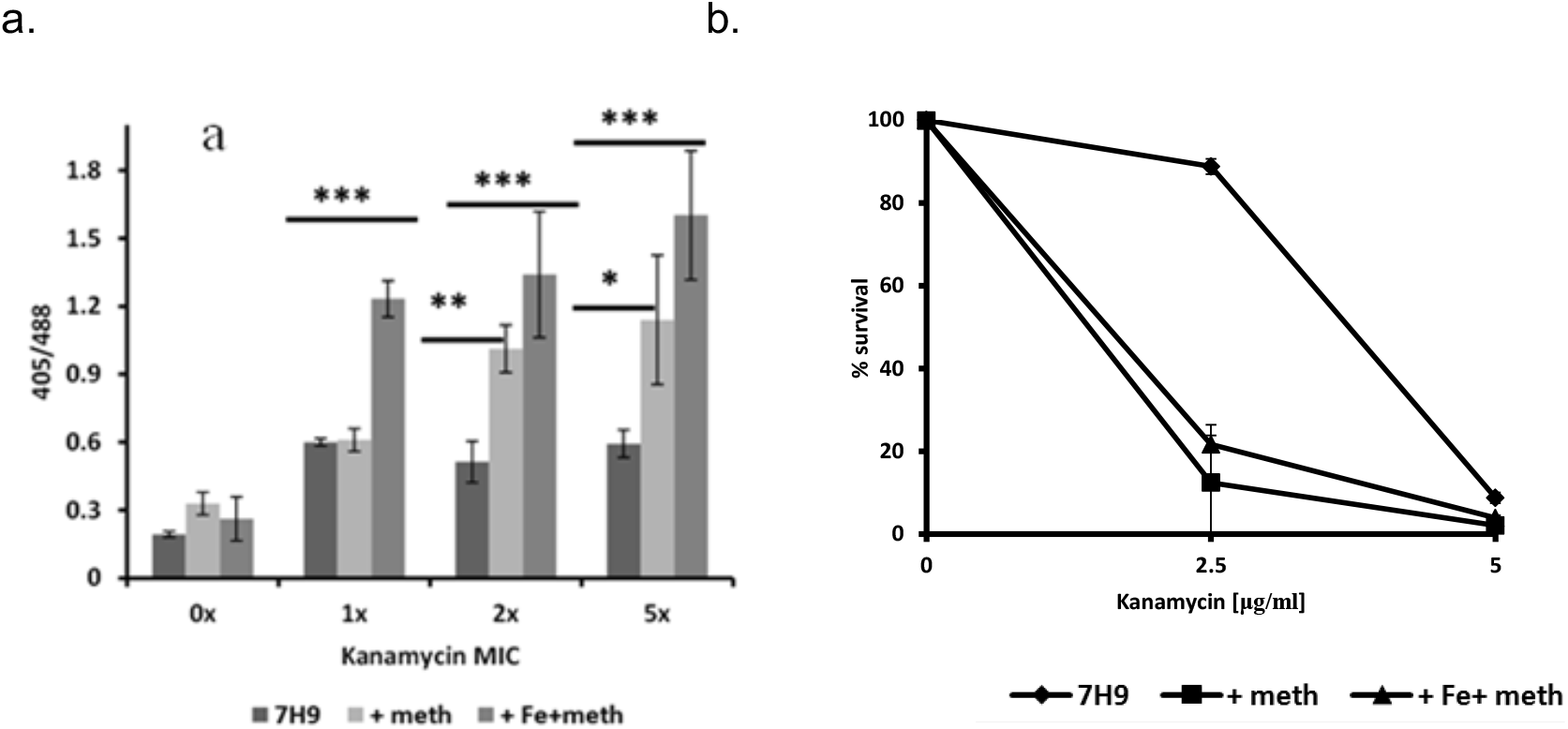
a) Oxidative stress induced by Kanamycin in presence of Fe and methionine. Statistical analysis was done using One way analysis of variance by Tukey’s test wherein the treatments were compared with control ***p<0.001; **p<0.01; b) Effect of exogenous methionine and Fe on kanamycin toxicity/tolerance in *Mtb* H37Rv

**Figure 5.**
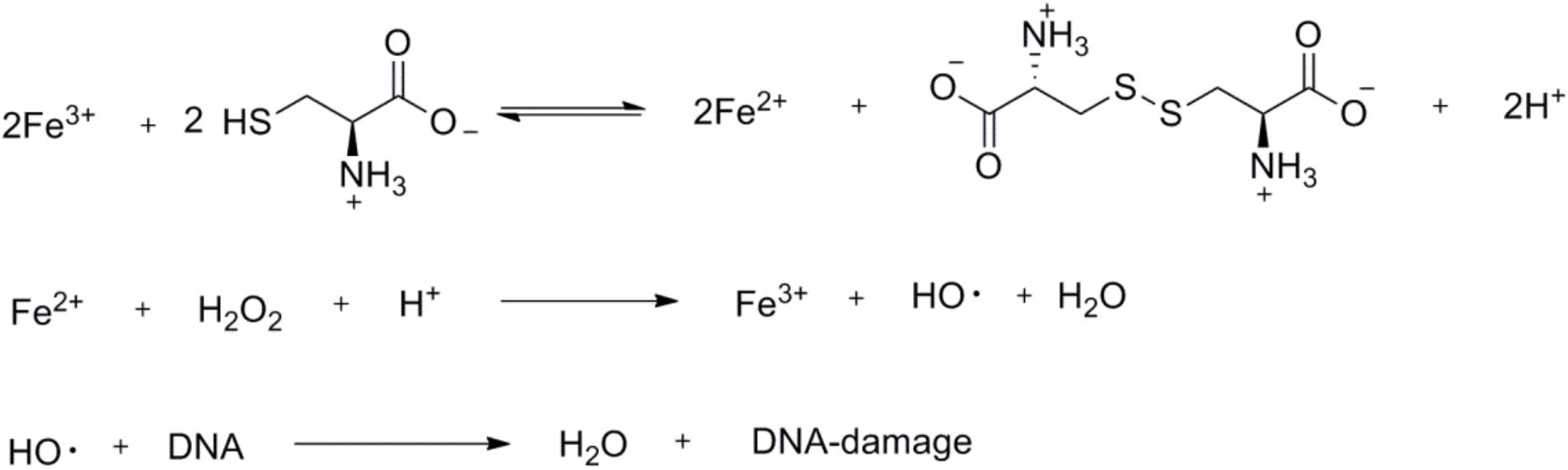
Cysteine mediated Fe+3 to Fe+2 and Fenton reaction.

## Discussion

The ability of mycobacterium to synthesize or acquire sulfur containing bio-molecules is central for its viability and defense against oxidative stress during phagocytosis. Mycobacteria can obtain sulfur by transporting sulfate via the sulfate transporter or transporting methionine via a methionine transporter, although the genes that encode the methionine transporter were not identified (11,15). However, biochemical studies indicate that methionine, but notcysteine, auxotrophs of *M. tuberculosis* could be isolated (16,17). In accordance with these observations, functional evidence for the existence of reverse transsulfuration, which utilizes methionine as sulfur source in mycobacteria, is reported and important methionine intermediates were characterized (11). Despite the functional and biochemical information about the reverse transsulfuration pathway, critical data about how methionine affects the levels of intracellular sulfur containing bio-molecules and how these changes affect intracellular redox potentials of mycobacterium in response to methionine stimulation is not clearly understood. To understand the intricacies of the thiol manipulation by mycobacteria, we have knocked out some important genes from mycobacteria and created mutants of respective gene using parish and storks method (18). In this present study we stimulated wild type and mutant mycobacteria with methionine and measured the intracellular redox potentials and quantified the relative levels of intracellular thiols using Mrx1-roGFP2(19) and monobromobimane (20,21) methods respectively.

Mrx1-roGFP2 is a non invasive tool based on genetically encoded redox sensitive fluorescent probes to perform real-time measurement of mycothiol redox potential (*E*_*MSH*_) in mycobacterium. This approach was utilized to measure the *E*_*MSH*_ of virulent and a virulent mycobacterial strains, including drug-resistant clinical isolates were successfully measured (19). Having this tool enabled us to measure *E*_*MSH*_ of wild type and sulfate assimilation pathway gene knockout strains of *M. smegmatis* in presence and in absence of methionine. When compared to wild type *M. smegmatis cys*A mutant exhibited oxidized *E*_*MSH*_ while the *cys*H mutant displayed reduced *E*_*MSH*_. In theory both the mutants are not capable of forming cysteine and mycothiol and hence we should actually see oxidized *E*_*MSH*_. But *cysH* mutant were unable to grow in absence of methionine and hence the cultures of *cysH* were supplemeted with 0.3mM methionine (22). The methionine grown culture were starved of methionine by washing and re-suspending the overnight grown cells at which time *E*_*MSH*_ was calculated. We assume that prior treatment of methionine might have up regulated the cellular thiol levels, giving rise to the observed reduced *E*_*MSH*_ for *cysH*. However Addition of exogenous methionine restored the intracellular *E*_*MSH*_ values comparable to the wt in both the mutants. The *E*_*MSH*_ of the *cys*Q remained unaffected and was identical to the wt *M. smegmatis* which is consistent with the role of *cysQ*, which catalyses the hydrolysis of 3’-phosphate of PAP and PAPS and have no direct relation with the reductive sulfate assimilation pathway. Also, in accordance with role of Mycothione reductase, the ΔMtr mutant exhibited oxidized *E*_*MSH*_ that showed no significant change in *E*_*MSH*_ upon addition of methionine.

Having measured the ambient *E*_*MSH*_ of these strains, we next analyzed the ability of these strains to dynamically change *E*_*MSH*_ in response to low concentrations of H_2_O_2_ (500 uM). Furthermore, we also thought of examining the influence of methionine on the dynamic changes in intracellular *E*_*MSH*_ upon addition of H_2_O_2_. Typically, wt *M. smegmatis* shows a faster and transient increase in the 405/488 ratio upon exposure to low concentrations of H_2_O_2_, indicating oxidative and anti-oxidative oscillations in intrabacterial *E*_MSH_(19). We observed that oxidative stress induces similar degree of roGFP2 oxidation inside wt, cysA, and cysQ strains (as indicated by a comparable increase in 405/488 ratio in all three strains). However, a striking difference was evident during the recovery phase. The cysA mutant showed extended oxidation (or delayed recovery), followed by cysQ, and wt *M.smegmatis*. Although this was expected in the case of cysA mutant (because of oxidized steady-state *E*_*MSH*_), delayed recovery was surprising in case of cysQ mutant which showed an initial steady-state *E*_*MSH*_ value similar to wt *M. smegmatis*. Next, we analyzed the effect of exogenous methionine on the dynamic response. Interestingly, we observed that addition of methionine significantly prolonged the oxidative burst in all three strains. However, in comparison to with wt *Msm*, the effect was similar in case of cysQ mutant and substantial in case of cysA mutant. We did not observe any recovery from oxidative stress in case of cysA+ methionine mutants during the 30 min time course of the assay. Based on these results we infer that addition of methionine may alter homeostasis of the intracellular thiol pool of the cells possibly leading to an increased production of free thiols which might have oxidised by the action of external hydrogen peroxide which might have lead to the oxidised states that we have observed in Figure 1. Similar observations were made in *E.coli* when cysteine homeostasis is disrupted, intracellular cysteine acts as an adventitious reductant of free iron and thereby promotes oxidative stress and oxidative DNA damage (12).

In order to address the delay in oxidative burst recovery that we observed, we next sought to profile intracellular thiols using monobromobimane coupled to liquid chromatography (20,21). We were surprised to see cysteine, cys-GlcN-Ins and mycothiol levels were found to be elevated 1 to 5 fold when cells were grown in presence of methionine. Cellular thiols are normally maintained in a highly reduced state. To determine the disulfide content we first reacted the thiols with *N*-ethylmaleimide, then reduced the soluble disulfides with dithiothreitol, and finally labeled the released thiols with monobromobimane for analysis by HPLC. This method cannot be used for CoA, because many acyl-CoA’s in the cell will undergo transacylation with dithiothreitol to release CoA not derived from disulfides(23). We found cystine, cys-GlcN-Ins and mycothiol disulfides levels were increased 1 to 5 fold when cultures were grown in presence of methionine. When *M. smegmatis* is grown in the absence of methionine the dominant thiol in *M. smegmatis* is MSH, followed by cysteine which is consistent with previous observation (13). During routine growth, cells maintain small cysteine pools, and cysteine is not a major contributor to DNA damage. Thus, the homeostatic control of cysteine is important in conferring resistance to oxidants. The elevated levels of intracellular thiols up on treatment with methionine have some unexpected consequences. The facile autoxidation of Cys has been attributed to the presence of a free amino group located beta to the thiol group that facilitate binding of heavy metals to catalyze thiol autoxidation. Cys-GlcN-Ins also possesses this functionality(13) and was observed to be oxidised considerably. In formyl-Cys-GlcN-Ins, succ-Cys-GlcN-Ins, and mycothiol, the amino group is acylated so that autoxidation is expected to be much slower which is consistent with the fact that mycobacterium can accomodate these thiols in higher levels without toxicity. However, in normal growth conditions slow auto oxidation of cellular thiols might also be coupled with the activity of mycothione reductase and thioredoxin to maintain optimal levels of free intracellular thiols. Biochemical studies show that both thiols (10^−3^ M) and NAD(P)H (10^−3^ M) (24,25) can transfer electrons to free iron in vitro and are far more abundant in vivo than is O^-2^. Free reduced flavins are also efficient reductants of iron, and in fact, their accumulation in respiration-inhibited cells causes a large increase in vulnerability to DNA damage (26-28). In Gram-positive and Gram-negative bacteria, a common mechanism of action by bactericidal drugs, regardless of their target, involves the generation of hydroxyl radicals by the Fenton reaction(29-31). Kinetic studies conducted by Jameson et al indicated that Fe^+3^ can be reduced to Fe^+2^ in presence of cysteine or other related thiols (32,33). The reduction of Fe^+3^ shown to occur through a complex formation between iron and cysteine followed by internal electron transfer (32-34).

Cysteine mediated Fe^+^3 to Fe^+2^ and Fenton reaction:

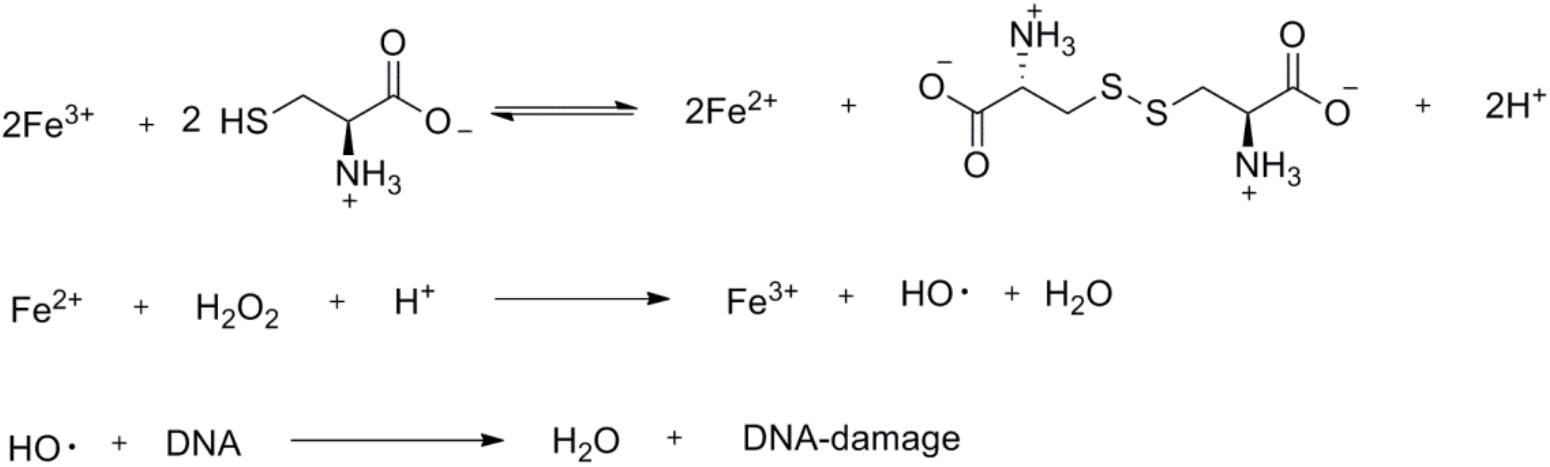

Taken together, the above data indicate the possible use of methionine and iron to elevate *in vivo* oxidative stress via Fenton reaction and can be augment with the anti-biotics by co-administering methionine and iron supplements. in line with this hypothesis it has been proposed that different classes of bactericidal antibiotics, regardless of their drug–target interactions, generate varying levels of deleterious reactive oxygen species (ROS) that contribute to cell killing (35,36). More recently vitamin c induced acceleration of the Fenton type reaction, which is responsible for increase in intracellular ROS, in Mycobacterium (29) is reported. In addition to these important observations, antibiotic induced redox related physiological changes were reported for several classes of antibiotics to treat several bacterial strains including mycobacterium (28,29,37-46). Based on these evidences we have conducted *in vitro* dose response studies of kanamycin in presence and in absence of methionine (0.3mM) and iron (0.85mM). Taken together the studies conducted here showed that the activity of kanamycin is found to be enhanced by approximately ten fold in presence of methionine and iron aganist *M. smegmatis*, indicating possible augmentation of the oxidative stress produced by antibiotic, methionine and iron treatment. Similar results were obtained when we challenged M. tuberculosis with kanamycin, methionine and iron. At MIC concentration kanamycin showed two fold incresed oxidative stress in presence of methionie and iron and cleared ninety percentof pathogens at 2.5μg/ml concentration. In contrast in the absence of methionie and iron at the same concentration ninety percent of pathogens are still viable. This observation is remarkable and could be easily translated to clinic. Both methionine and iron are being used in clinical settings as supplements and are inexpensive. Hence formulation of kanamycin doses in presence of those supplements would be highly beneficial for controlling the multi drug resistant tuberculosis.

Clinical management of tuberculosis involves use of first line drugs isoniazid, INH and rifampicin, which are bactericidal and can kill 99% of mycobacterium tuberculosis cells in vitro within a week. Unfortunately, resistance emerges rapidly for these antibiotics both *in vitro* and *in vivo* (47,48) and mismanagement of MDR-TB may turn in to more fatal XDR-TB. Hence, novel drugs that effectively kill mycobacterium quickly through a unique mode of action are urgently needed to address the global rise of multi drug resistant pathogens. However finding a new drug is not easy and consume significant amount of time and resources. In order to address this issue as an alternate way, in this present study we report methionine treatment causes a temporary loss of cysteine homeostasis, with intracellular thiol pools increasing about five fold in mycobacterium, could reduces ferric iron with exceptional speed and produce reactive oxygen radicals. This surge in oxidative stress was coupled with kanamycin activity and obtained 2 fold improved activity of the antibiotic against Mycobacterium tuberculosis. During routine growth, cells maintain small cysteine pools, and at that low concentration cysteine is not a contributor to DNA damage(12). Thus, the homeostatic control of cysteine levels is important in conferring resistance to oxidants. In our studies, the enhancement of bactericidal activity of methionine supplemented antibiotics against *M. tuberculosis*has been shown to be dependent on free thiol concentrations, intracellular iron, and associated ROS production through fenton type reaction. This study enlightens the possible benefits of adding methionine and iron to an antituberculosis regimen that use kanamycin as second line injectable drug to treat MDR-TB.

## Materials and Methods

### Generation of M. smegmatis, MC^2^155, gene knockout strains

Using the two-step counter selection strategy described by Parish and Stoker (49), the gene knockout vectors were constructed by first inserting a hygromycin resistance cassette into p2NIL. Second, 2-kb regions upstream and downstream of respective genes, *cysH, cysQ*, mycothione reductase (*MTR*) and *cysHQ*, were amplified from *M. smegmatis*, MC^2^155, DNA and cloned into the multicloning site flanking the hygromycin resistance gene. Finally, the *lacZ*-*sacB* counter-selection cassette from pGOAL17 was inserted. The Vector DNAs were pretreated with 100 mJ UV light cm^-2^ and electroporated into separate *M. smegmatis* cell samples. Single crossovers were selected on 7H11 plates with 40 μg/ml 5-bromo4-chloro-3-indolyl β-D-galactoside (X-Gal), 50 μg/ml hygromycin, and 20 μg/ml kanamycin. Double crossovers were selected on 7H11 plates containing 40 μg/ml X-Gal, 2% sucrose, and 50 μg/ml hygromycin. The candidate double crossover mutants were screened by PCR and then confirmed by Southern blot hybridization.

### Estimation of intracellular Mycothiol and its major components using HPLC and bromobimane

The assay method for estimation of thiols and disulfides were adopted from the previously published methods by Newton and Fahey *et al*. Cells from Mycobacterium smegmatis MC^2^155 wild type and gene knockpout strains were cultured in Middlebrook 7H9 medium supplemented with 10% ADC. Cultures were grown at 30 degress in 50 mls of medium in 100ml Ehrlynmeyer flasks shaking at 200 rpm. Cultures were allowed to grow to OD_600_ 0.5 and then challenged with methionine and allowed to grow another hour to approximately OD_600_ 0.68. ***For thiol analysis***: Cells, 10ml, were harvested by centrifugation and re-suspended in 0.5 ml warmed 50% acetonitrile -20mM Hepes buffer pH 8.0. Monobromobimane was added to 2mM and cells were incubated at 60 °C for 15 min with occasional vortexing to assure cell lysis and labelling of mycothiol and its components. Cell debris were removed by centrifugation and samples were suitably diluted in 10mM HCL. Samples were injected onto a C18, 4.6×250mm Beckman ultra sphere column set to 390 nm excitation and 475 nM emission. The column was equilibrated with buffer A, 0.25 M acetic acid pH 3.6, and eluted with buffer B, HPLC grade methanol, with a gradient of 0 to10 min-10%, 15 min-18% buffer B, 30 min-27% buffer B, 32 min-100% buffer B, 34 min-10%B. Data was analyzed using the software provided in the Agilent chem station data analysis module. ***For disulfide analysis*:** Cells, 10ml, were harvested by centrifugation and re-suspended in 0.5 ml warmed 50% acetonitrile -20mM Hepes buffer pH 8.0. *N*-ethylmaleimide (NEM) was added to 5mM and cells were incubated at 60 °C for 15 min with occasional vortexing to assure cell lysis and labelling of free thiols. Cooled the samples on ice and clarified the supernatant by centrifugation at 13000xg for 5 min and removed 100μL sample to separate tube and reacted with 2mM bromobimane for 10 min at 60 °C to serve as control sample to identify the fluorescent peaks not derived from thiols. To the remaining 900μL added 2-mercaptoethanol to a final concentration of 5mM, to scavenge the excess NEM, and incubated the samples at room temperature for 10 min and concentrated the samples using ultrahigh vaccum at room temperature. To 100μL of the concentrate added 2mM DTT at room temperature and reduced the disulfides for 15 min. The samples were acidified by adding 2μL of 5M methanesulfonicacid and suitably diluted with 10mM methanesulfonicacid prior to HPLC analysis. Samples were injected onto a C18, 4.6×250mm Beckman ultra sphere column set to 390 nm excitation and 475 nM emission. The column was equilibrated with buffer A, 0.25 M acetic acid pH 3.6, and eluted with buffer B, HPLC grade methanol, with a gradient of 0 to10 min-10%, 15 min-18% buffer B, 30 min-27% buffer B, 32 min-100% buffer B, 34 min-10%B. Data was analyzed using the software provided in the Agilent chem station data analysis module. Standard peaks for the reported thiols were assigned based on the previous published retention times for the respective thiols (50).

### Effect of exogenous methionine and Fe on kanamycin toxicity in M. smegmatis MC^2^155

A well defined colony of wild type *M. Smegmatis* was selected and carefully transferred to 3ml of 7H9 media with ADC supplements in a sterile culture tube. After 36 hours of incubation at 37 °C obtained fresh culture of growing bacteria. The fresh culture was used to inoculate the secondary culture in 1: 10000 dilution in 7H9 media. The cultures were incubated while shaking in preence and absence of methionine and iron for 36 hours. The IC_50_ was determined as the concentration of Kanamycin that prevented 50% of growth of *M. smegmatis*.

### M. smegmatis killing experiments and DNA toxicity experiments using Kanamycin Measurement of M. smegmatis DNA damage by quantitative PCR (qPCR) assays

Total genomic DNA was isolated from 10 ml of culture using the procedure described in supporting information section. The extracted DNA was quantified using nanodrop spectrophotometer. For primers, 10-kb fragments of adenosine 5’-phosphosulfate reductase regions were used. Primer sequences were as follows: 5’-ATGAGCGGCGAGACAAC-CAGGCTGACCGAACCGCAA (forward primer) and 5’-TCACGAGGCGTGCAACCCG-CATTCGGTCTTGG (reverse primer). PCR was performed using whole genome amplification kit (New England biolabs). The 50μL PCR mixture contained 0.05 to 0.5 ng of genomic DNA as a template, a 300 nM concentration (each) of the two primers, 350 μM each of the four deoxynucleoside triphosphates provided along the kit, 10X PCR buffer with 1.75 mM MgCl2, and 1 μL of DNA polymerase mix. Thermal cycling was performed with Eppendorf PCR thermal cycler. The genomic DNA was initially denatured for 1 min at 94°C, and then the DNA was subjected to 25 cycles of PCR, with 1cycle consisting of denaturation at 94°C for 15 s and annealing and extension at 68°C for 12 min. A final extension step at 72°C was performed for 10 min at the completion of the profile. PCR products were separated by 1% agarose gel electrophoresis, stained with ethidium bromide, scanned with Alpha innotech FluorchemQ imager, and the DNA bands were quantified with band analysis tool provided in the software of the instrument.

## Figures and Tables

**Table 1:** E_MSH_ measurement in *M. Smegmatis* sulfate assimilation pathway knock out strains.

| Strains | $E_{MSH}$ (mV) | |
| --- | --- | --- |
|  | -methionine | 0.3mM methionine |
| **WT** | -298.5 ± 0.5 | -297 |
| **cysA** | -281.17 ± 2.78 | -290 ± 0.29 |
| **cysQ** | -298 ± 2.03 | -301 ± 1.80 |
| **cysH<sup>1</sup>** | -311 ± 1.7 | -292 ± 3.2 |
| **Mtr** | -277.4 ± 0.41 | -274 ± 0.96 |

<sup>1</sup> Since cysH mutants didn't grow without methionine therefore the cells were grown in 7H9 media supplemented with 0.3 mM methionine. The methionine grown culture was starved of methionine by washing and resuspending the overnight grown cells ( $OD_{600} = 2.0$ ) in fresh 7H9 media supplemented with ADS. This was followed by incubation at 37°C for 2h at which time $E_{MSH}$ was calculated.

## Supporting information

To isolate genomic DNA of *M. smegmatis:* A fresh colony from the overnight incubating 7H10+OADC plate were isolated and inoculated to 7H9 ADC media and grown them overnight. The cells were harvested and the cell pellet was collected.

*Lysis of cells:* The collected pellets were re suspended in 0.6ml of TE (50mM Tris, 25mM EDTA) pH 7.4 buffer and added 400μL of 50mg/ml of lysozyme solution. Added 4μl of RNase Solution (Promega) to the cell lysate. Inverted the tube 2–5 times to mix. the cells were Incubated at 37°C for 16 h and mixed occasionally by gentle agitation.

*Removal of protein and cellular debris:* To the suspended cells added 150 μL of 10 mg/mL proteinase K and mixed gently and incubated at 45°C for 16h. The samples were cooled to room temperature and added 0.5 mL of phenol/chloroform/isopropanol (1:1:1) and mixed gently by inverting tube five times. Repeated the inversion steps four times over 30 min. *(Phenol was saturated with TE buffer pH 8.0 (equal volumes of phenol and TE buffer was shaken vigorously and allowed to stand at room temperature to get the phase separation). This step eliminates the toxic free radicals that exist in the phenol. These free radicals if allowed to pass in to the DNA purification procedure they will cause DNA strand breaks and we may not see the right size or results of DNA on agarose gels)*. Then the samples were centrifuged at 3000*g* at room temperature for 20 min. The protein precipitated was sedimented in between the organic and aqueous layers and removed the upper aqueous phase without disturbing the protein precipitate. To the aqueous phase collected added 0.5 mL of chloroform/isopropanol (1:1), mixed gently and centrifuged at 3000*g* at room temperature for 30 min. The upper aqueous layer was separated. To the recovered aqueous layer added 2μL of pellet paint dye (EMD bioscience) and added 0.1 volume of 3M sodium acetate pH 5.2 solution and mixed. To the mixed solution added 1.5 volumes of isopropanol and inverted the tube for 5 to 6 times, white floccules of DNA were obtained and it seems that the DNA floccules were slowly growing back to the solution. Then the solution was kept incubating at -20°C for 30 mins. *(Adding pellet paint is necessary for visualizing DNA. Trials without adding pellet paint was resulted in transparent DNA precipitate which cannot be detected using visual eye. Also, DNA precipitation was tried using equal volumes of isopropanol but was resulted in limited success. Thus the isopropanol volume was adjusted to 1.5 olumes of the aqueous phase that is recovered from step 9.)* Then the incubated solution was centrifuged at 3000g for 30 min at 4°C. A pink color precipitate of DNA was appeared at the bottom of the eppendorf tube. All of the supernatant liquor was pipetted without disturbing the bottom DNA. The bottom DNA was washed with 70%ethanol and dried. The DNA obtained was solubilised in 100μL of TE buffer.

